# Directed evolution predicts cytochrome *b* G37V target site modification as probable adaptive mechanism towards the QiI fungicide fenpicoxamid in *Zymoseptoria tritici*

**DOI:** 10.1101/2021.09.03.458847

**Authors:** Guillaume Fouché, Thomas Michel, Anaïs Lalève, Nick X Wang, David H Young, Brigitte Meunier, Danièle Debieu, Sabine Fillinger, Anne-Sophie Walker

## Abstract

Acquired resistance is a threat for antifungal efficacy in medicine and agriculture. The diversity of possible resistance mechanisms, as well as the highly adaptive traits of pathogens make it difficult to predict evolutionary outcomes of treatments. We used directed evolution as an approach to assess the risk of resistance to the new fungicide fenpicoxamid in the wheat pathogenic fungus *Zymoseptoria tritici*. Fenpicoxamid inhibits complexIII of the respiratory chain at the ubiquinone reduction site (Qi site) of the mitochondrially encoded cytochrome b, a different site than the widely-used strobilurins which the respiratory complex by binding to the ubiquinol oxidation site (Q_o_ site). We identified the G37V change, within the cytochrome *b* Q_i_ site, as the most likely resistance mechanism to be selected in *Z. tritici*. This change triggered high fenpicoxamid resistance and halved the enzymatic activity of cytochrome *b*, despite no significant penalty for *in vitro* growth. In addition, we identified a negative cross-resistance between isolates harboring G37V or G143A, a Q_o_ site change previously selected by strobilurins. Moreover, double mutants were less resistant to both QiIs and QoIs compared to single mutants. This work is a proof of concept that experimental evolution can be used to predict adaptation to fungicides, and provides new perspectives for the management of QiIs.

**Originality-Significance Statement:**

- The highly adaptive traits of pathogens render evolutionary outcomes of antifungal treatments difficult to predict.
- We used directed evolution to assess the risk of resistance to the new fungicide fenpicoxamid in the wheat pathogenic fungus *Zymoseptoria tritici*.
- We identified a target modification as the most likely resistance mechanism to be selected.
- This change triggered high fenpicoxamid resistance and halved the activity of the target enzyme despite no significant penalty for *in vitro* growth.
- This work supports the use of experimental evolution as a method to predict adaptation to fungicides and provides important information for the management of QiIs.

## INTRODUCTION

Acquired resistance is a phenotypic adaptation, mainly in pathogens, pests, weeds or cancer cells, in response to drug selection pressure (Hawkins *et al*., 2019). Resistance is responsible for major economic losses and public health concerns in both agriculture and medicine (Ahmad and Khan, 2019; Maragakis *et al*., 2008). Therefore, predicting the adaptation to new active ingredients and novel modes of action is a critical significance for implementing anti-resistance strategies at the onset of selection. In practice, resistance predictions are complicated by the diversity of resistance mechanisms found in nature, and the particular interactions between active ingredients, and the population and biology of targeted organisms. Resistance selection in the laboratory can be achieved by two different approaches. The classical method involves random mutagenesis, and searching for the resistance-conferring mutation(s). This approach has proven effective in medicine to understand resistance mechanisms in cancer cells (Azam *et al*., 2003) or pathogenic yeasts (Rawal *et al*., 2013), and in agriculture to explore insecticide (McKenzie and Batterham, 1998) or fungicide resistance (Hawkins and Fraaije, 2016; Scalliet *et al*., 2012). However, random mutagenesis achieved using UV or chemical exposure may cause multiple mutations including rearrangements throughout the genome and make it difficult to detect and correlate the resistance phenotype with a specific mutation. Moreover, some mutations recovered *in vitro* may never occur in nature under selection pressure from agrochemicals or drugs (Hawkins and Fraaije, 2016). The use of molecular techniques to perform site-directed mutagenesis avoids additional mutations, but focuses only on single mutations involving the biochemical target, with no information on other possible mechanisms. A second approach is directed evolution, which consists in allowing strains or cell lines to evolve under drug selection pressure, but without mutagenic agents, to accelerate resistance selection in miniaturized conditions (for review, Kawecki *et al*., 2012). This method mimics natural selection, and may select any putative resistance mechanism. In particular, it has been used with cancer cells (Kern and Weisenthal, 1990), bacteria (Jahn *et al*., 2017; Orencia *et al*., 2001), nematodes (Lopes *et al*., 2008) and fungi (Cowen *et al*., 2001; Gutiérrez-Alonso *et al*., 2017; Schoustra *et al*., 2006). However, this approach is conducted without the host and is therefore not ideal either. Thus, the best prediction relies on the combination of different approaches that would provide converging data.

In agriculture, fungal pathogens represent the biggest threat for crop production (Savary *et al*., 2019), and there is a continual need for new fungicides due to resistance. As key organelles, fungal mitochondria represent a target of choice for fungicide development. Seven different fungicidal modes of action used in agriculture target the mitochondrial respiratory chain (https://www.r4p-inra.fr/fr). Although known mainly for their role in energy production, mitochondria also participate in many other metabolic processes and govern the life and death cycle of eukaryotic cells (reviewed in van der Bliek *et al*., 2017). In eukaryotes, most cellular ATP results from the activity of the mitochondrial oxidative phosphorylation (OXPHOS) system. Five multi-subunit protein complexes embedded in the inner mitochondrial membrane (IMM) form the OXPHOS system. Complexes I to IV transfer electrons to oxygen and produce energy, in the form of a transmembrane electrochemical gradient of protons, that is used by complex V (or ATP synthase) for the production of ATP (reviewed in van der Bliek *et al*., 2017). Some organisms also possess alternative oxidases (AOX), which allow ATP production while bypassing complexes III and IV (reviewed in Day *et al*., 1995). Complex II inhibitors, known as SDHIs (succinate dehydrogenase inhibitors) and complex III quinone outside inhibitors (QoIs, or strobilurins), are the most widely used respiratory inhibitor fungicides in agriculture. SDHI resistance has been reported in many fungi, including *Aspergillus flavus* (Masiello *et al*., 2020), *Botrytis cinerea* (Fernández-Ortuño *et al*., 2017), *Sclerotinia sclerotiorum* (Peng *et al*., 2020), *Alternaria alternata* (Fan *et al*., 2015) and *Zymoseptoria tritici* (Dooley *et al*., 2016; Steinhauer *et al*., 2019). Prior to the commercial use of SDHIs to control *Z. tritici*, mutagenesis studies in the laboratory identified some mutations which later occurred in the field, but also produced mutations which have not yet been found in nature (Fraaije *et al*., 2012; Rehfus *et al*., 2018; Scalliet *et al*., 2012). In the case of QoIs, their repeated and unrestricted use in the field (Bartlett *et al*., 2002) quickly triggered the emergence of target site resistance in many fungal pathogens. This rapid emergence and spread occurred in multiple species (Hawkins and Fraaije, 2021), and had not been predicted. The most common resistance mutation leads to the G143A change, and has been selected in *Z. tritici* (Fraaije *et al*., 2005) and repeatedly in many other pathogens (Hawkins and Fraaije, 2021). In *Z. tritici*, resistance is now generalized in populations from Western Europe (Garnault *et al*., 2019; Kildea *et al*., 2019).

Complex III (or cytochrome *bc*_1_ complex) catalyzes the transfer of electrons from ubiquinol to cytochrome *c*, and couples this electron transfer to the translocation of protons across the IMM (reviewed in Brzezinski *et al*., 2021). The complex operates as a dimer. Its monomeric unit consists of 10 or 11 different polypeptides. Three subunits, cytochrome *b*, cytochrome *c*_1_ and the Rieske iron-sulphur protein, form the catalytic core of the enzyme. Cytochrome *b* is encoded by the mitochondrial genome in all eukaryotes, and the other subunits are encoded by nuclear genes (reviewed in Berry *et al*., 2000; Crofts, 2004; Fisher *et al*., 2020). The complex possesses two distinct ubiquinone-binding sites situated on opposite sides of the IMM; the ubiquinol oxidation site (Q_o_ site) located towards the IMM outer side, and the ubiquinone reduction site (Q_i_ site) located towards the inner, or matrix, side. The Q_o_ and Q_i_ sites are both in the membrane-spanning cytochrome *b* protein (Hunte *et al*., 2000). Fungicide inhibitors of complex III bind to either the Q_o_ site (QoIs) or the Q_i_ site (QiIs), although the oomycete fungicide ametoctradin appears to bind at both sites (reviewed in Fisher *et al*., 2020). Two QiIs, cyazofamid and amisulbrom, are currently used to control Oomycete diseases. The picolinamide compound fenpicoxamid (Inatreq™ Active, Trademark of Corteva Agriscience and its affiliated companies) is the first QiI fungicide active against ascomycete pathogens and will provide a new mode of action in the cereal market to control *Z. tritici* and other diseases (Owen *et al*., 2017). Derived from the natural antifungal compound UK-2A (Ueki et al., 1996), fenpicoxamid is rapidly metabolized in fungal or wheat cells back into UK-2A, which is responsible for its fungicidal activity (Owen *et al*., 2017).

*Z. tritici* is the causal agent of wheat *Septoria tritici* blotch (STB). STB is the economically most important wheat disease in Europe, in terms of potential yield losses and cost of disease control (Fones and Gurr, 2015; Savary *et al*., 2019). Due to its high genomic plasticity, ability to perform sexual reproduction and to spread over long distances through ascospore production, *Z. tritici* populations have developed resistance towards all currently used unisite modes of action (Garnault *et al*., 2019). This highlights the relevance of resistance risk for new fungicides targeting *Z. tritici* such as fenpicoxamid.

Since QiI resistance has been described for cyazofamid and amisulbrom in Oomycetes and because respiration inhibitors are generally at risk, we anticipate that resistance might also affect fenpicoxamid. Different resistance mechanisms can be envisaged in *Z. tritici* (Fig. 1). Firstly, QiIs are subject to resistance risk due to target modification as reported in *Plasmopara viticola* (Fontaine *et al*., 2019), and in several human protozoan parasites (reviewed in Mounkoro *et al*., 2019). Mutations conferring resistance to UK-2A and the structurally related natural product antimycin A were also identified in the model organism *Saccharomyces cerevisiae*. In addition, the selection pressure-driven maternal behavior of *Z. tritici* field isolates during sexual reproduction has been reported for QoIs (Kema *et al*., 2018). This behavior might accelerate the spread of resistance mechanisms transferred through the mitochondrial genome.

**Fig. 1:**
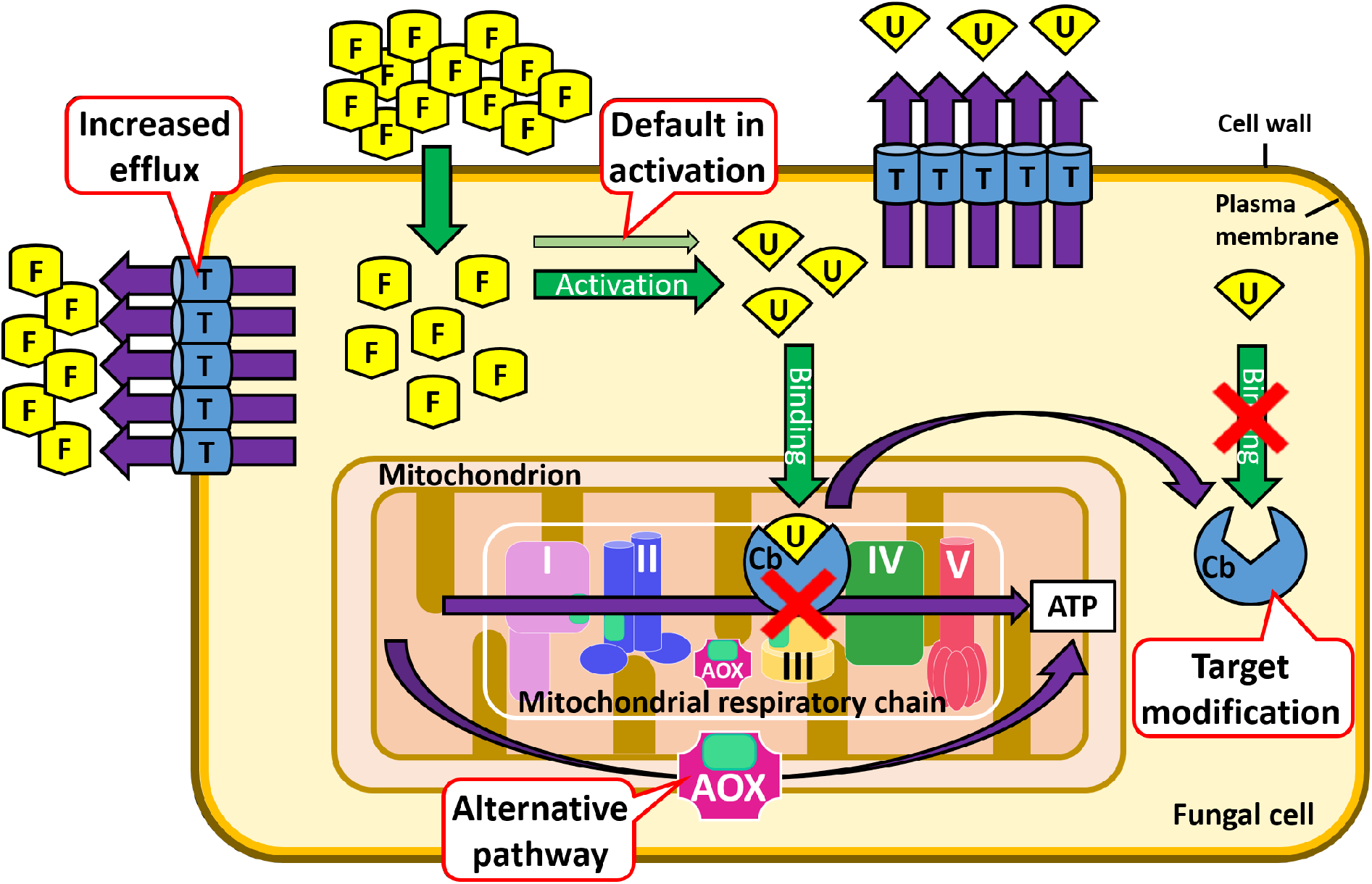
Potential resistance mechanisms which might reduce susceptibility of *Z. tritici* field isolates to fenpicoxamid: target modification, increased efflux, alternative pathway and default in activation. **F:** fenpicoxamid; **U:** UK-2A; **Cb:** cytochrome *b*; **AOX:** alternative oxidase; **T:** membrane transporter.

Increased efflux in *Z. triciti* can lead to reduced sensitivity to almost all unisite fungicides, including mitochondrial respiration inhibitors (Leroux and Walker, 2011; Omrane *et al*., 2017). This phenotype, called MDR for Multi-Drug Resistance, is associated with overexpression of the membrane transporter gene *MFS1* (Omrane *et al*., 2015). Another potential resistance mechanism, applicable to both QiIs and QoIs, is activation of the AOX pathway. This has been observed in some *P. viticola* strains, which are resistant to amisulbrom and ametoctradin but contain no mutation in the *CYTb* gene (Fontaine *et al*., 2019). Finally, a default in the activation process by which fenpicoxamid is converted to UK-2A could contribute to reduced sensitivity even though such type of resistance is rarely observed in fungi (e.g., Tellier *et al*., 2009).

Because of its biology, and ease of handling in liquid culture due to its yeast-like form (Francisco *et al*., 2019; Steinberg, 2015), *Z. tritici* is a convenient organism to predict adaptation to a fungicide. The goal of our study is to assess the risk of resistance to fenpicoxamid by using experimental evolution to produce resistant mutants in *Z. tritici*. Putative resistance mechanisms in isolated mutants were evaluated, revealing a target site modification, which was analyzed for its effect on complex III activity and growth rate as indicators of fitness, and on binding of UK-2A using molecular docking analysis.

## RESULTS

### Impact of enhanced efflux on *Z. tritici* sensitivity to fenpicoxamid

Because increased efflux causes low to moderate resistance to current respiration inhibitors (*i*.*e*. SDHIs and QoIs; Leroux and Walker, 2011) and other fungicides in *Z. tritici*, we assumed that this resistance mechanism might affect fenpicoxamid. Moreover, MDR strains have been detected in Western European populations at increasing frequencies each year, and represent roughly 40% of the French population (Garnault *et al*., 2019). We first evaluated the impact of enhanced efflux on fenpicoxamid susceptibility by determining EC_50_ values for inhibition of apical germ-tube elongation in solid medium, and *in vitro* growth in liquid cultures (Table 1), for representative MDR isolates. Efflux derived MDR leads in both assays to around 10-fold increased EC_50_ values reflecting low to moderate resistance levels. This in the same range as the resistance factors measured for different modes of action (Omrane *et al*., 2017).

**Table 1.** Impact of increased efflux on sensitivity to fenpicoxamid in *Z. tritici*.

| Assay | Isolate | *MFS1*-<br>genotype<sup>a</sup> | Cytochrome *b* | EC<sub>50</sub><sup>b</sup> (µg/mL) | RF |
| --- | --- | --- | --- | --- | --- |
| Germ-tube<br>elongation<br>(48h) | IPO323 | WT | WT | 0.004 ± 0.002 | - |
|  | 14-H-K3 | WT | G143A | 0.008 ± 0.002 | - |
|  | 09-ASA-3a-PZ | MDR<sup>Type I</sup> | G143A | 0.027 ± 0.006 | 3<sup>c</sup> |
|  | 09-CB1 | MDR<sup>Type I</sup> | G143A | 0.054 ± 0.023 | 7<sup>c</sup> |
| *In vitro*<br>growth<br>(3 days) | IPO323 | WT | WT | 0.004 ± 0.002 | - |
|  | 35F22 | WT | WT | 0.003 ± 0.001 | - |
|  | 35F6 | WT | G143A | 0.0020 ± 0.0007 | - |
|  | 35G5 | WT | G143A | 0.0014 ± 0.0006 | - |
|  | 37-15 | WT | G143A | 0.0018 ± 0.0004 | - |
|  | 37-16 | WT | G143A | 0.0009 ± 0.0002 | - |
|  | 11183 | MDR<sup>Type I</sup> | WT | 0.0293 ± 0.0006 | 8<sup>d</sup> |
|  | 35C3 | MDR<sup>Type I</sup> | G143A | 0.017 ± 0.004 | 11<sup>c</sup> |
|  | 35C4 | MDR<sup>Type I</sup> | G143A | 0.019 ± 0.003 | 12<sup>c</sup> |
|  | 37-59 | MDR<sup>Type I</sup> | G143A | 0.019 ± 0.005 | 12<sup>c</sup> |

<sup>a</sup>*MFS1* genotype nomenclature according to Omrane *et al.*, 2017.
<sup>b</sup>EC<sub>50</sub> is the fungicide concentration inhibiting 50% of apical germ tube elongation or growth, compared to untreated control. Each EC<sub>50</sub> value is the average of at least three independent assays.
<sup>c</sup>RF is the ratio between the EC<sub>50</sub> of the MDR isolate and the EC<sub>50</sub> value or the average of EC<sub>50</sub> values from the G143A isolates.
<sup>d</sup>RF is the ratio between the EC<sub>50</sub> of the MDR isolate and the average of EC<sub>50</sub> values from the WT isolates.

### Selection of fenpicoxamid-resistant mutants

Experimental evolution speeds up resistance selection under miniaturized laboratory conditions. Resistance to fenpicoxamid was selected using directed evolution in eleven *Z. tritici* ancestral isolates exhibiting different genetic backgrounds. Isolates used were the reference strain IPO323 (Goodwin *et al*., 2011), as well as MDR, QoI-resistant (StrR) and MDR+StrR strains. These last three genotypes are the most frequent ones in European populations (Garnault *et al*., 2019). Strains were cultivated in liquid medium amended with fenpicoxamid at the minimal inhibitory concentration (MIC) or at 25 times the MIC (25MIC), over eight cycles (see Experimental procedures for details). At the end of each cycle, spores were plated on fenpicoxamid containing selection medium to isolate resistant colonies. The experiments allowed the selection of almost 400 isolates which produced colony-like growth on fenpicoxamid-containing medium (Fig. 2). However, after subculturing even once on fenpicoxamid-free medium and/or storage, 88% of these isolates lost fenpicoxamid resistance. This transient resistance phenomenon was observed for isolates derived from all ancestral strains, regardless of the genetic background (Fig. 2). Stable resistant isolates were only derived from IPO323 (WT) and 37-16 (StrR).

**Fig. 2:**
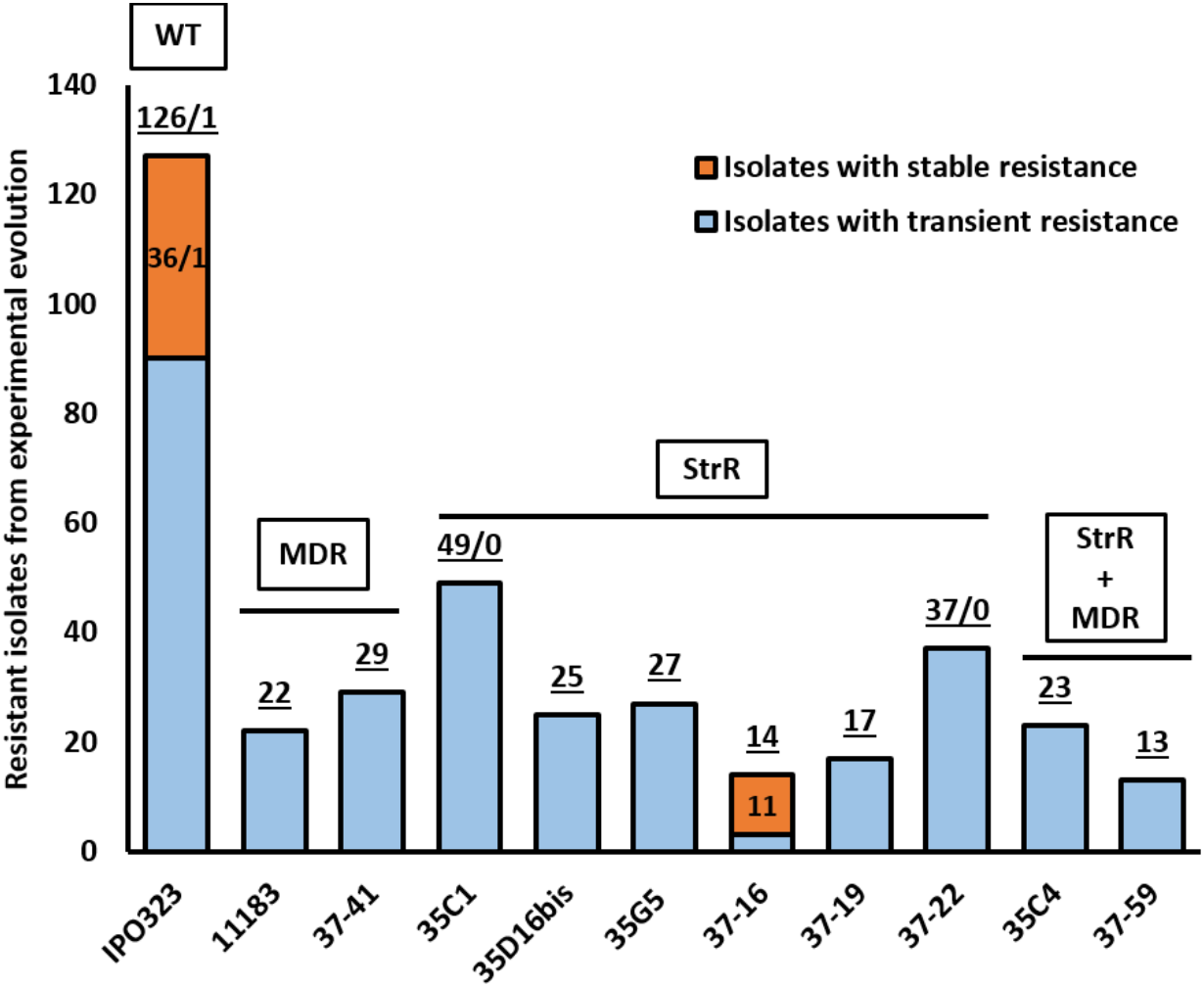
Number of fenpicoxamid-resistant isolates generated by experimental evolution for each ancestral strain. Total number of isolates is underlined. When relevant, the first number represents isolates selected at the MIC, and the second number represents isolates selected at 25MIC for concerned ancestral strains. Phenotypes of ancestral strains are specified in boxes.

The sensitive IPO323 ancestral strain generated 127 resistant isolates in total, among which 37 (29%) were stable. Individuals with stable resistance were isolated in 50% of the independently evolved IPO323 lines. However, only one isolate with stable resistance was selected in lines evolving at 25MIC, while all others were selected at the MIC. MDR strains, regardless of their cytochrome *b* genotype, never provided stable resistant isolates. Resistance to fenpicoxamid was also explored for six different StrR strains; only one strain (37-16) yielded isolates with stable resistance, representing one line out of 25 (4%).

### Characterization of fenpicoxamid-resistant mutants

Resistant isolates were characterized for their putative resistance mechanisms using a droplet test on fungicide-amended YPD medium (Fig. 3). Growth on fenpicoxamid and UK-2A were both assessed, in order to detect a potential impairment in fenpicoxamid metabolic activation in the resistant isolates. Azoxystrobin sensitivity was tested to identify possible cross-resistance between QoIs and QiIs, which would be expected from overexpression of AOX. Growth on fenpicoxamid in the presence of AOX inhibitors, SHAM or propyl gallate, was also assessed to screen for this resistance mechanism, and provided a means of evaluating this mechanism in the StrR isolates where assessment of cross-resistance to azoxystrobin was not applicable. Finally, tolnaftate sensitivity was used to discriminate isolates exhibiting enhanced efflux leading to MDR in several fungal species (Kretschmer *et al*., 2009; Leroux and Walker, 2011; Leroux *et al*., 2013; Nakaune *et al*., 1998; Omrane *et al*., 2017; Sang *et al*., 2015).

**Fig. 3:**
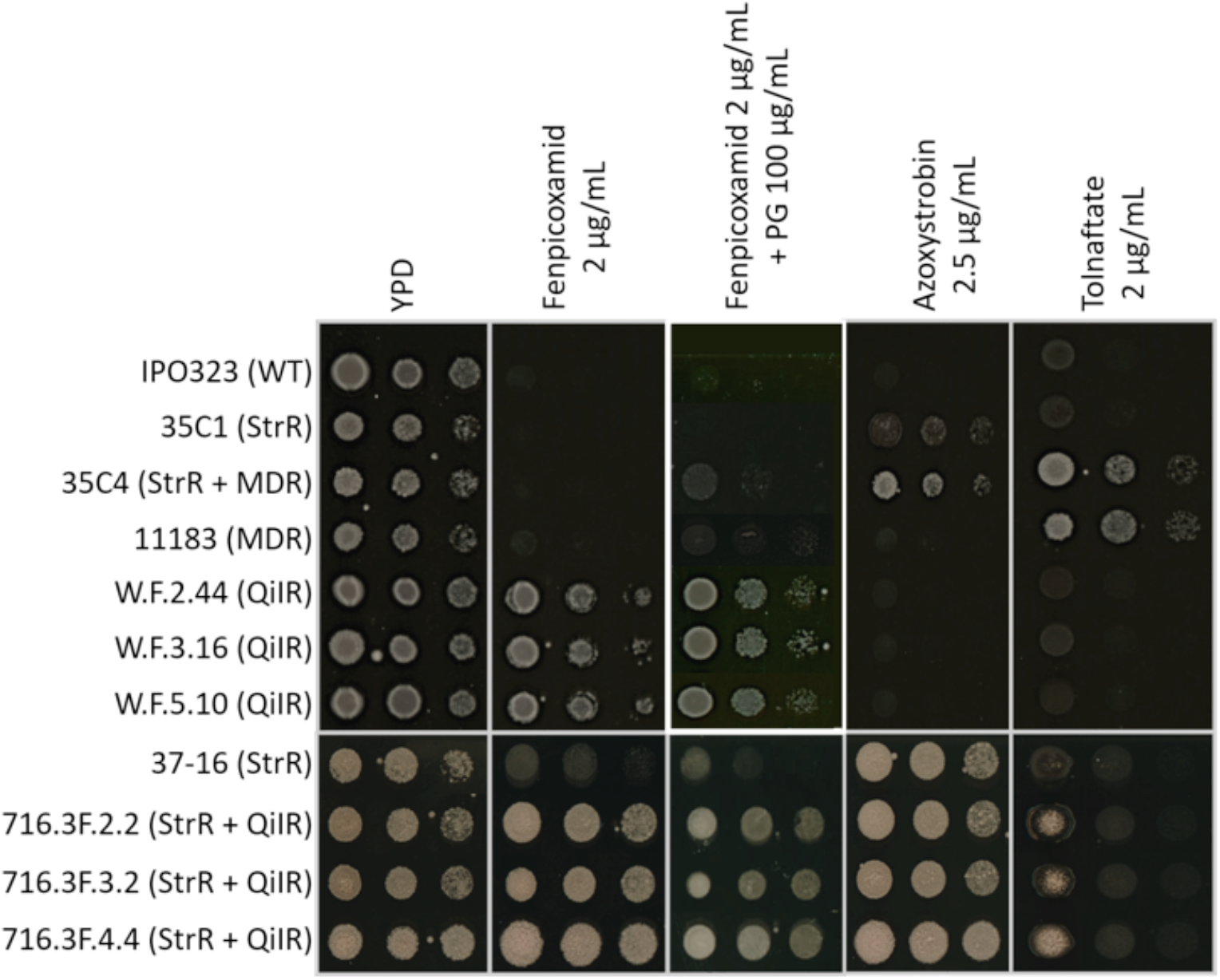
Phenotype of representative resistant isolates obtained by experimental evolution, in comparison to their ancestral strains. The isolates W.F.2.44, W.F.3.16 and W.F.5.10 were derived from IPO323, while isolates 716.3F.2.2, 716.3F.3.2 and 716.3F.4.4 were derived from 37-16 (StrR).

All evolved isolates displayed growth on fenpicoxamid at 2 µg/mL (its approximate solubility limit in aqueous media), a concentration which prevented growth of all ancestral strains (Fig. 3). Those isolates also grew on 0.5 µg/mL UK-2A-amended YPD (data not shown), excluding a default in fenpicoxamid activation as the resistance mechanism. As expected, IPO323-derived isolates remained susceptible to the QoI azoxystrobin, while isolates derived from 37-16, bearing the cytochrome *b* G143A change, were resistant (Fig. 3). In addition, growth on fenpicoxamid was not prevented by the addition of 100 µg/mL of the AOX inhibitors propyl gallate (Fig. 3) or SHAM (data not shown), for any of the resistant isolates, demonstrating that AOX overexpression was not involved in resistance. None of these isolates was able to grow on 2 µg/mL tolnaftate (Fig. 3), ruling out increased efflux as a resistance mechanism in the selected isolates.

After phenotypic analysis, the cytochrome *b* encoding gene (*CYTb*) was sequenced for 29 isolates. All sequences exhibited as sole *CYTb* mutation the same G to T substitution at codon 37 (GGA to GTA), leading to replacement of the glycine residue by valine. This suppports target site modification as the most plausible resistance mechanism towards fenpicoxamid.

### Consequences of the G37V change on QiI and QoI susceptibility during *in vitro* growth

We first assessed the resistance to fenpicoxamid associated with target modification at the fungal cell level. Isolates bearing only the G37V change were highly resistant to UK-2A (Table 2), with EC_50_ values ranging from 0.8 to 1.6 µg/mL, and RFs higher than 200. The double mutants, displaying both the G37V and the G143A changes, were also highly resistant to UK-2A, with EC_50_ values ranging from 0.28 to 1.28 µg/mL, and RFs ranging from 79 to 356 when compared to IPO323. Relative to G143A ancestral isolates, RFs for the double G37V+G143A mutants ranged from 183 to 828.

**Table 2.** Susceptibility to UK-2A and azoxystrobin as measured *via in vitro* growth of *Z. tritici* isolates.

| Isolate | Cytochrome *b* | UK-2A |  |  | Azoxystrobin |  |  |
| --- | --- | --- | --- | --- | --- | --- | --- |
|  |  | EC<sub>50</sub> (µg/mL)* | RF<sub>WT</sub><sup>a</sup> | RF<sub>StrR</sub><sup>b</sup> | EC<sub>50</sub> (µg/mL) | RF<sub>WT</sub><sup>a</sup> | RF<sub>StrR</sub><sup>b</sup> |
| IPO323 | WT | 0.004 ± 0.002 | - | 3 | 0.043 ± 0.022 | - | > 0.01 |
| 35F22 | WT | 0.003 ± 0.001 | - | 2 | 0.035 ± 0.012 | - | > 0.01 |
| W.F.2.44 | G37V | 0.80 ± 0.18 | 222 | 516 | 0.026 ± 0.009 | 0.7 | > 0.01 |
| W.F.4.14 | G37V | 1.0 ± 0.4 | 286 | 665 | 0.019 ± 0.01 | 0.5 | > 0.01 |
| W.F.5.10 | G37V | 1.6 ± 0.9 | 451 | 1050 | 0.022 ± 0.014 | 0.6 | > 0.01 |
| 35F6 | G143A | 0.0020 ± 0.0007 | 0.6 | - | 5.1 ± 1.8 | 132 | - |
| 35G5 | G143A | 0.0014 ± 0.0006 | 0.4 | - | 3.5 ± 2.3 | 92 | - |
| 37-15 | G143A | 0.0018 ± 0.0004 | 0.5 | - | 6.5 ± 2.1 | 168 | - |
| 37-16 | G143A | 0.0009 ± 0.0002 | 0.3 | - | 4.1 ± 1.5 | 106 | - |
| 716.3F.2.2 | G37V+G143A | 0.5 ± 0.1 | 140 | 327 | 3.0 ± 0.2 | 78 | 0.6 |
| 716.3F.2.3 | G37V+G143A | 0.75 ± 0.09 | 209 | 486 | 3.6 ± 0.4 | 93 | 0.7 |
| 716.3F.2.4 | G37V+G143A | 1.28 ± 0.08 | 356 | 828 | 3.2 ± 0.1 | 83 | 0.7 |
| 716.3F.3.1 | G37V+G143A | 0.8 ± 0.05 | 223 | 519 | 5.3 ± 0.3 | 138 | 1 |
| 716.3F.3.2 | G37V+G143A | 0.28 ± 0.09 | 79 | 183 | 2.0 ± 0.5 | 53 | 0.4 |
| 716.3F.4.1 | G37V+G143A | 0.46 ± 0.07 | 127 | 296 | 1.7 ± 0.5 | 45 | 0.4 |
| 716.3F.4.2 | G37V+G143A | 0.81 ± 0.09 | 226 | 526 | 2.2 ± 0.2 | 58 | 0.5 |
| 716.3F.4.3 | G37V+G143A | 0.66 ± 0.09 | 185 | 430 | 0.7 ± 0.2 | 19 | 0.1 |
| 716.3F.4.4 | G37V+G143A | 0.73 ± 0.39 | 203 | 473 | 2.9 ± 0.7 | 76 | 0.6 |
| 716.3F.4.5 | G37V+G143A | 0.94 ± 0.33 | 261 | 608 | 2.6 ± 0.3 | 66 | 0.5 |

<sup>a</sup>RF<sub>WT</sub> is the ratio between the EC<sub>50</sub> of the resistant isolate and the average of EC<sub>50</sub> values from the WT isolates. Each EC<sub>50</sub> value is the average of at least three independent assays.
<sup>b</sup>RF<sub>StrR</sub> is the ratio between the EC<sub>50</sub> of the resistant isolate and the average of EC<sub>50</sub> values from the G143A isolates. Each EC<sub>50</sub> value is the average of at least three independent assays.
\*On average, EC<sub>50</sub> of UK-2A were 1.15 µg/mL for G37V isolates, 0.0015 µg/mL for G143A isolates, and 0.72 µg/mL for double mutants; EC<sub>50</sub> of azoxystrobin were 0.022 µg/mL for G37V isolates, 4.79 µg/mL for G143A isolates, and 2.73 µg/mL for double mutants.

Resistance to QoIs was also measured for double mutants, using azoxystrobin. RFs ranged from 10 to 138 when compared to IPO323. However, RFs of the G37V+G143A double mutants when compared to G143A single mutants were below 1, demonstrating that their resistance towards QoIs was reduced.

Comparing the RFs to QoIs and QiIs of the G37V and G143A single mutants, respectively, we observed that RFs were systematically lower than one (Table 2); the G37V mutants were up to two-times more susceptible to QoIs than the WT, and the G143A mutants up to three times more susceptible to QiIs than the WT. These results suggest increased susceptibility of cytochrome *b* mutants to inhibitors which act at the opposite binding site, with QiIR isolates being more sensitive to QoIs and StrR isolates more sensitive to QiIs.

### Consequences of the G37V change on complex III (or cytochrome *bc*_1_ complex) activity and fungal growth rate

To evaluate the impact of G37V on enzyme activity, we first measured the activity of complex III without fungicide inhibition (Fig. 4). Activity was assessed spectrophotometrically by monitoring cytochrome *c* reduction. It was normalized by the independent enzyme activity of complex IV, as complex III concentrations might vary among the different mitochondrial preparations. The G37V change reduced complex III activity in the mutant strains; for IPO323-derived mutants activity was reduced by approximately 50%, and for mutants derived from the strobilurin-resistant parent 37-16 activity was reduced by about 30%. The G143A change also appeared to cause a slight (20%) decrease in complex III activity (Fig. 4) based on comparison of activity for IPO323 and 37-16, however the difference is within the range of variability for this type of experiment.

**Fig. 4:**
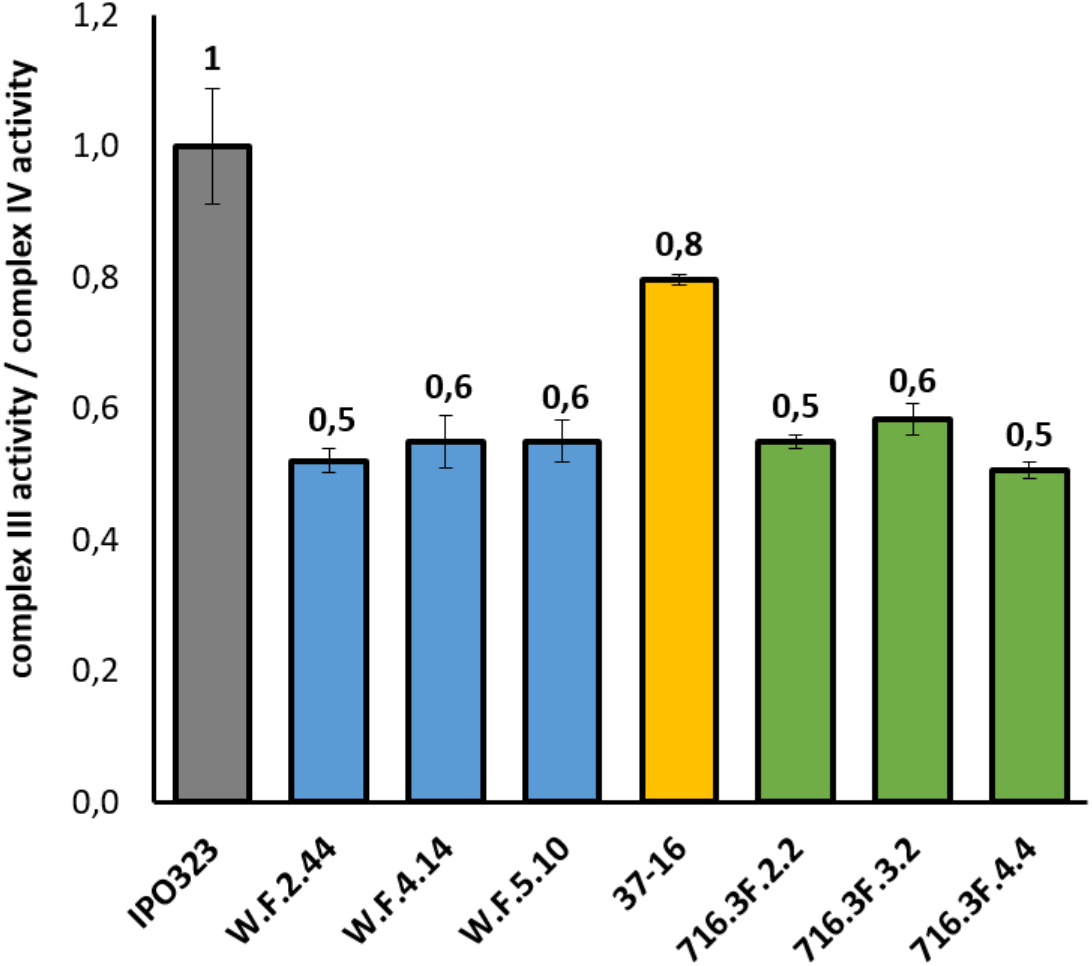
Complex III activity of different *Z. tritici* isolates. For each mitochondrial sample, complex III activity was measured as the rate of cytochrome *c* reduction, and normalized by complex IV activity. Each assay was repeated at least three times, and the values averaged. Values were normalized to a constant ratio of 1 for IPO323.

Although complex III activity was impaired by the G37V change, the growth rate in YPD-amended microtiter plates of fenpicoxamid-resistant mutants was similar relative to their ancestral strains when measured over 60 hours (Fig. 5).

**Fig. 5:**
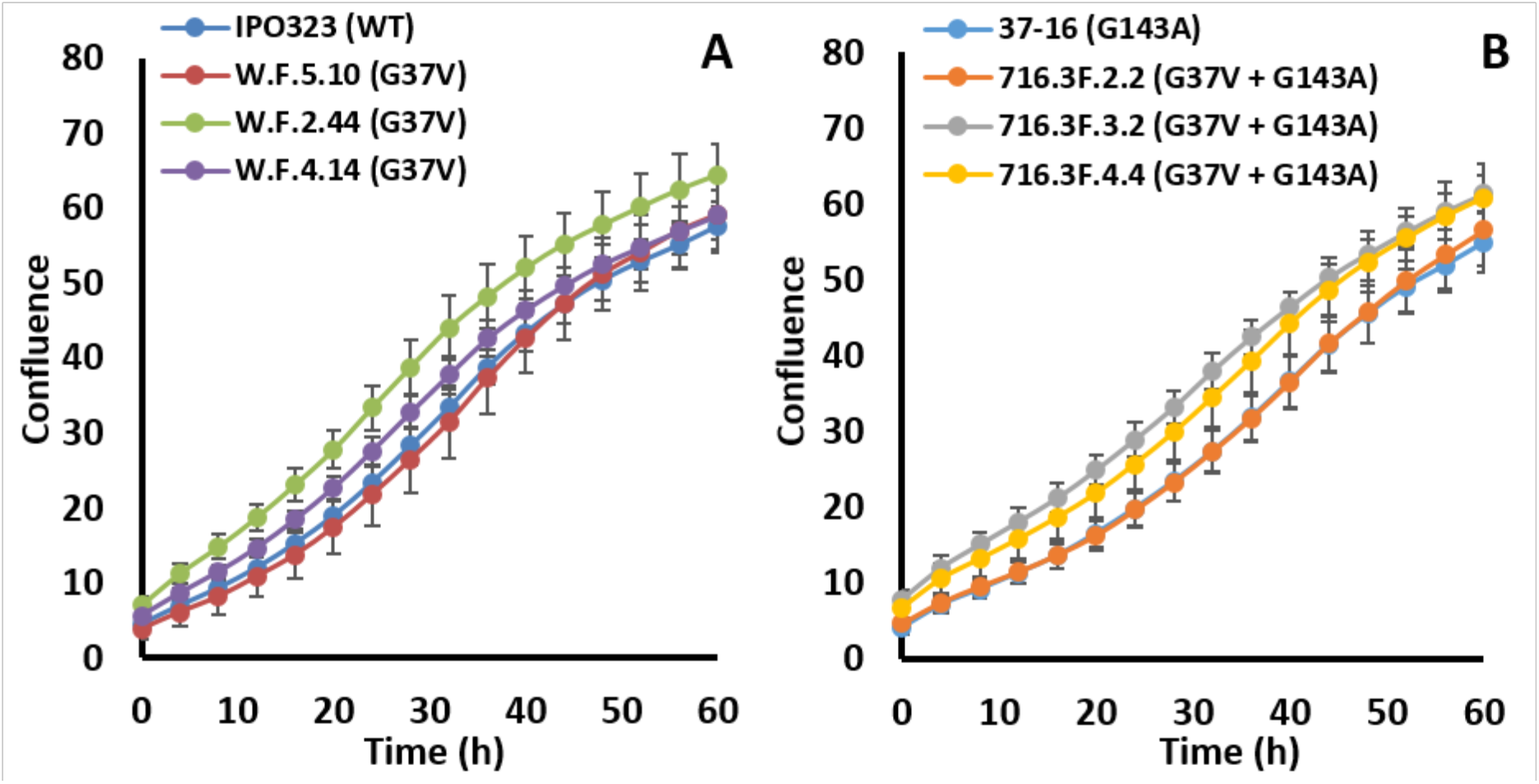
Growth of G37V mutants compared to their ancestral strains. **A**. Growth of G37V single mutants compared to IPO323. **B**. Growth of G37V+G143A double mutants compared to the G143A strain 37-16. The maximum growth rates estimated after using a logistic regression model were not significantly different between IPO323 and the single mutants. For the double mutants, the maximum growth rate of 716.3F.3.2 was not significantly different from that of 37-16, and the other two double mutants showed a very slight apparent increase in growth rate. However, growth rates of the 3 mutants were not statistically different from each other, and clearly any differences relative to 37-16 were small.

To confirm that the G37V modification of cytochrome *b* conferred resistance by reducing enzyme sensitivity to UK-2A, I_50_ values were determined using crude mitochondrial extracts from the parent and mutant strains (Table 3). RFs based on I_50_ values for UK-2A ranged from 81 to 171 for the cytochrome *b* G37V mutants, relative to their ancestral IPO323 strain. In contrast, RFs for cytochrome *b* G37V+G143A mutants ranged from 9 to 15 when compared to IPO323 and from 16 to 26 when compared to their G143A parent strain 37-16. Therefore, double mutant enzymes were about eleven times more sensitive to UK-2A than the enzymes of G37V single mutants. Complex III of the G143A parent strain 37-16 was also 1.6 fold more susceptible to UK-2A than the enzyme from IPO323.

**Table 3.** Impact of substitutions on complex III sensitivity to UK-2A and azoxystrobin

| Isolate | Cytochrome *b* | UK-2A |  |  | Azoxystrobin |  |
| --- | --- | --- | --- | --- | --- | --- |
|  |  | I<sub>50</sub> (μM) | RF<sup>a</sup> | RF<sup>b</sup> | I<sub>50</sub> (μM) | RF<sup>a</sup> |
| IPO323 | WT | 0.012 ± 0.004 | - | 1.7 | 0.113 ± 0.003 | - |
| W.F.2.44 | G37V | 2.1 ± 0.5 | 171 | 290 | 0.090 ± 0.004 | 0.5 |
| W.F.4.14 | G37V | 0.968 ± 0.005 | 81 | 136 | 0.10 ± 0.03 | 0.6 |
| W.F.5.10 | G37V | 1.8 ± 0.1 | 151 | 256 | 0.10 ± 0.01 | 0.6 |
| 37-16 | G143A | 0.0071 ± 0.0004 | 0.6 | - | > 15 | > 100 |
| 716.3F.2.2 | G37V+G143A | 0.18 ± 0.01 | 15 | 26 | > 15 | > 100 |
| 716.3F.3.2 | G37V+G143A | 0.114 ± 0.001 | 9 | 16 | > 15 | > 100 |
| 716.3F.4.4 | G37V+G143A | 0.140 ± 0.001 | 12 | 20 | > 15 | > 100 |

<sup>a</sup>RF is the ratio between the I<sub>50</sub> value of the resistant mutant and the I<sub>50</sub> value of the WT IPO323 strain.
<sup>b</sup>RF is the ratio between the I<sub>50</sub> value of the resistant mutant and the I<sub>50</sub> value of the G143A parent strain 37-16.

Regarding sensitivity to azoxystrobin, complex III from the G37V single mutants was two times more susceptible than the enzyme from their parent strain. Enzyme RFs for strains with the G143A change could not be precisely determined because the I_50_ values were above the azoxystrobin solubility limit in the aqueous assay buffer. Therefore, the precise impact of the additional G37V change on sensitivity to azoxystrobin could not be assessed in the G143A background.

### Effect of the G37V change on binding of UK-2A at the cytochrome *b* Q_i_ site

A *Z. tritici* homology model was built based on the crystal structure of the *S. cerevisiae* cytochrome *bc*_1_ complex. Docking of UK-2A in the *Z*.*tritici* Q_i_ site showed the importance of cytochrome *b* residue D230 (D229 in *S. cerevisiae*, Fig. 6) to stabilize the binding pose, as well as the proximity of G37 to the exocyclic methyl group and the ester tail on the bislactone ring of UK-2A. Figure 6 shows how the substitution of G37 by a valine, with its large apolar isopropyl side chain, approximately halves the distance between UK-2A and residue 37.

**Fig. 6:**
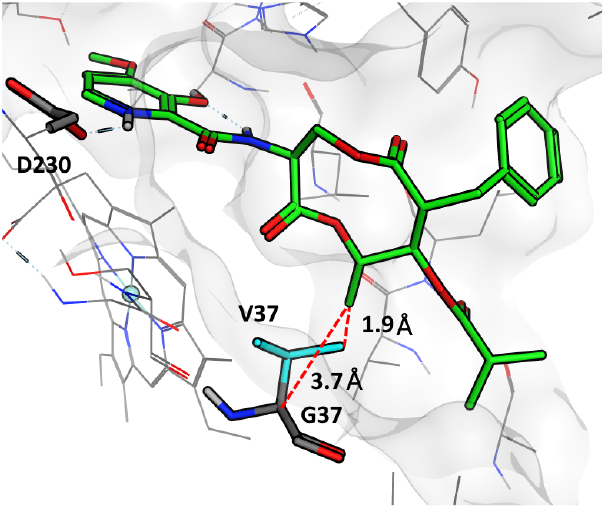
Binding of UK-2A at the Q_i_ site of *Z. tritici* cytochrome *b*. Consequences of the G37V substitution are shown by the reduction of the distance (red dotted lines) between the amino acid in position 37 and the UK-2A exocyclic methyl group.

## DISCUSSION

Fenpicoxamid is a new fungicide, developed to control *Z. tritici*, the causal agent of wheat STB. It inhibits the cytochrome *bc*_1_ complex by binding to the Q_i_ site, which represents a new mode of action for *Z. tritici* control. Predicting adaptation to antifungal compounds in fungal populations represents a challenge for evolutionary biology, with practical implications in agriculture and medicine for the management of pathogens. The resistance of *Z. tritici* to fenpicoxamid constitutes a relevant model to study these questions, as outlined in the introduction.

We used directed evolution to select *Z. tritici* isolates resistant to fenpicoxamid, as successfully used for other modes of action (Hawkins and Fraaije, 2016). Evolved isolates displaying a stable phenotype all harbored the G37V change in the Q_i_ site of cytochrome *b*. This change led to fenpicoxamid resistance at both the whole cell and enzyme levels. However, the G37V resistance mutation led to a 50% reduction of complex III activity, despite no measurable impact on *in vitro* growth rate. This change also increased the susceptibility of resistant isolates to Q_o_ site inhibitors. Conversely, the G143A change at the Q_o_ site of cytochrome *b* increased sensitivity towards fenpicoxamid. In addition, RFs towards QoIs and QiIs in the G37V+G143A double mutants were on average lower than for single mutants. These findings may be related in some way to the smaller proportion of G37V isolates selected in G143A backgrounds, as compared to the WT isolate. However, a larger set of directed evolution data from experiments using multiple ancestral strains with WT, G143A-cytochrome *b* and MDR backgrounds would be required to test rigorously the hypothesis that G37V mutants may be selected less readily in a G143A background, and further explore the potential link to observed negative cross-resistance between QoIs and QiIs. Altogether, directed evolution suggests that target alteration due to the G37V change is a likely resistance mechanism to emerge in *Z. tritici* populations under fenpicoxamid selection pressure.

### Directed evolution as a tool to predict resistance selection

Among different evolutionary pathways which could allow adaptation to fenpicoxamid in *Z. tritici*, the most viable should be determined by the relative fitness of selected mutants, *i*.*e*. their capacity to outcompete other strains in the selecting environment. In our experiments, directed evolution was used to mimic and accelerate the processes of mutation and natural selection in naive ancestral strains. Resistant isolates were selected as early as the second cycle of the experiment, confirming the utility of this approach. Our protocol did not use a mutagenic agent to make directed evolution as realistic as possible in simulating natural selection. Natural selection depends on random genetic changes introduced by errors during replication or mutation repair, and even the most beneficial mutations can be lost due to genetic drift, especially in small populations subjected to environmental factors which restrict multiplication and spread (Baer *et al*., 2007; Liu and Zhang, 2021). A similar situation can be considered to occur in experimental evolution experiments due to the small number of cells transferred in each cycle. This may explain why not all lines derived from the same ancestral isolate led to resistance. We also noticed variations according to the ancestral isolate, possibly due to epistasis and/or insufficient fitness. For example, stable resistance was never selected in MDR ancestors, and found less often in G143A strains. In the latter case, this may reflect a stronger fitness penalty for isolates displaying alterations in both the Q_i_ and Q_o_ sites of cytochrome *b*. The ability of epistatic interactions between mutations in OXPHOS complexes to increase fitness penalties has also been described in *B. cinerea* field isolates which display target site based resistance to both QoIs and SDHIs (Veloukas *et al*., 2014). Most striking in our study was the repeated selection of G37V, rather than other changes in cytochrome *b*, in multiple independent lines and different genetic backgrounds, suggesting an optimal trade-off between resistance and fitness penalty for this substitution. In our experiments, additional mutations may have been selected in some isolates as reflected in the variability of RFs observed for both the enzyme activity (81 to 171 and 9 to 15 for G37V and G37V+G143A, respectively) and *in vitro* growth (222 to 451 and 79 to 356). G37V was selected much more readily in lines exposed to the MIC as selection pressure, as compared to lines exposed to 25MIC. This is consistent with evidence from multiple organisms that sub-lethal doses enable mutations to emerge more readily (Amaradasa and Everhart, 2016; Andersson and Hughes, 2014; Busi and Powles, 2009; Rix and Cutler, 2018). Also, this highlights the importance of fungicide concentration in the experimental evolution approach.

### Target site resistance as the most probable resistance mechanism

We proposed four potential resistance mechanisms that might affect susceptibility to fenpicoxamid in *Z. tritici*. Reduced conversion of fenpicoxamid to UK-2A could be envisaged as a potential resistance mechanism. A default in metabolic activation is a known resistance mechanism for the human antifungal drug 5-flucytosine (Vermes *et al*., 2000) and is associated with natural insensitivity of some *Botrytis cinerea* strains to cymoxanil (Tellier *et al*., 2009). However, the scarcity of marketed active ingredients requiring activation, together with poor exploration, may explain why this mechanism is barely described. Resistance due to a default in activation was not detected in our experiments, as shown by the positive cross-resistance between fenpicoxamid and UK-2A in isolated mutants. By analogy to other masked ester molecules, the activation mechanism for fenpicoxamid is expected to involve carboxylesterase enzymes (Tian *et al*., 2012). Carboxylesterase(s) mediating the hydrolysis of fenpicoxamid into UK-2A have not been characterized. However, the ubiquitous occurrence of these enzymes and the fact that activation also operates *in planta* (Owen *et al*., 2017), suggests that this resistance mechanism is not the most likely to affect fenpicoxamid susceptibility.

A second potential resistance mechanism involves activation of the AOX-mediated alternative respiratory pathway. Some weakly resistant isolates derived from MDR ancestral strains displayed a slightly reduced growth on fenpicoxamid in the droplet test in the presence of the AOX inhibitor propyl gallate, suggesting that in some strains AOX overexpression could lead to a slight decrease in susceptibility to QiIs, especially if combined with MDR. Sequencing of the *CYTb* gene in these strains did not reveal any mutation, and expression levels were not explored to validate AOX involvement. AOX expression should affect QiI fungicides as well as QoIs. In this respect, it is of interest that some QoI-resistant isolates from Ireland displayed lower RFs in the presence of SHAM, suggesting that AOX expression levels may vary among individuals and could play a role in resistance to strobilurin fungicides (Kildea *et al*., 2019), although this effect is clearly minor compared to the impact of the G143A change. In addition, reliance on AOX is associated with lower energy production which may be insufficient to satisfy a high energy requirement during early stages of infection in *Z. tritici* (Miguez *et al*., 2004). Altogether, these findings suggest that AOX overexpression is unlikely to be a major resistance mechanism for fenpicoxamid in *Z. tritici* field populations.

Enhanced efflux, or multidrug resistance, is a general mechanism associated with positive cross resistance between DMIs, QoIs and SDHIs (Leroux and Walker, 2011). *In vitro* sensitivity to fenpicoxamid was reduced slightly in MDR field isolates (Table 1), and RFs for fenpicoxamid were similar to those for QoIs and SDHIs (RFs 2-15) (Leroux and Walker, 2011; Omrane *et al*., 2015, 2017). In our experiments, MDR was not detected in evolved mutants. A possible explanation is that the selection pressure was limited against fenpicoxamid since MDR is selected more readily in field populations sprayed with mixtures of fungicides with different modes of action (Garnault *et al*., unpublished). The frequency of MDR has increased each year in French populations since its first detection in 2008, and is generalized in some locations (Garnault *et al*., 2019). The effect of enhanced efflux alone on fenpicoxamid sensitivity is likely to be small. Nevertheless, it may insidiously contribute to increase RFs when combined with other resistance mechanisms, as it is largely installed in field populations and because sexual reproduction occurs multiple times per season in *Z. tritici* (Zhan *et al*., 2003).

Surprisingly, directed evolution also led to the selection of unexpected mechanisms associated with fenpicoxamid resistance. A high number of isolates, initially characterized as resistant, rapidly lost their resistance to fenpicoxamid when subcultured without fungicide. This phenomenon has already been described for QoI-resistant *Z. tritici* field isolates (Miguez *et al*., 2004). Miguez and colleagues isolated 46 QoI-resistant isolates growing on high azoxystrobin concentrations. However, only six isolates were still able to grow on azoxystrobin-amended medium after being subcultured on fungicide-free medium. This transient resistant state might correspond to “tolerance”, a phenomenon regularly observed in human pathogenic yeasts (reviewed in Berman and Krysan, 2020). Tolerance refers to a state in which susceptible isolates become able to grow in the presence of high antifungal concentrations, due to the modification of several metabolic pathways. Although it is a well-studied phenomenon in human pathogenic fungi, tolerance has been largely ignored in resistance studies concerning agricultural fungicides. A second hypothesis could be the existence of an unstable heteroplasmic state for mitochondrial mutations in *Z. tritici*, as already observed in other phytopathogenic fungi (Ishii *et al*., 2007, 2009). Heteroplasmy is defined as a cellular state where at least two different mitochondrial populations (or mitotypes) coexist in the same cell, and may be transient (Mendoza *et al*., 2020). In the case of reduced enzyme activity in resistant mitochondria, the parental mitotype would be preferentially selected, and fungal cells would rapidly return to homoplasmy in the absence of selection pressure (Christie *et al*., 2015; Ni *et al*., 2011). Whatever the underlying mechanism, these observations should prompt further investigation of transient resistant states induced by agricultural fungicides. It could be argued that these metabolic states may allow isolates to survive longer when exposed to QiI fungicides, therefore increasing the risk of selecting resistance mutations. On the other hand, if stable resistance requires further selection to achieve a homoplasmic state, the development of resistance may be delayed.

Alteration of the target site is the most common resistance mechanism in fungi (Lucas *et al*., 2015), and is therefore likely to be of greatest importance for fenpicoxamid resistance. Indeed, the G37V change in cytochrome *b* was repeatedly selected in independent lines and genetic backgrounds in our directed evolution studies and was associated with high RFs. In addition, mutations at this locus have conferred resistance to diverse QiIs in multiple organisms, as reviewed in Mounkoro *et al*., 2019. As examples, the G37A and G37V changes have been reported in lines of the human parasite *P. falciparum* resistant to the macrocyclic QiIs ML238 and BRD6923, and to antimycin A (Lukens *et al*., 2015). The G37C and G37V (di Rago and Colson, 1988) changes were associated with antimycin A resistance in the model organism *S. cerevisiae*, G37D and G37S resulted in ilicicolin H resistance (Ding *et al*., 2006), and the G37C change conferred resistance to UK-2A (Young *et al*., 2018). Moreover, the G37V change was also selected in *Z. tritici*, after repeated exposure to high concentrations of antimycin A (Fehr *et al*., 2016). Parallel evolution is frequent in fungicide resistance (Hawkins *et al*., 2019). A review of literature suggests that lab mutations associated with high RFs, and those found in multiple species, are more likely to be reported in the field (Hawkins and Fraaije, 2016, 2021). Altogether, our findings make the selection of G37V a realistic and most probable evolutionary scenario for *Z. tritici* populations exposed to fenpicoxamid selection pressure in the field, and suggest this could occur in both QoI-sensitive and resistant populations. This conclusion is also supported by the lack of an obvious fitness penalty on *in vitro* growth as measured in G37V single and G37V+G143A double mutants. Nevertheless, it will be important to assess fitness *in planta* during the infection cycle.

### G37V influences complex III activity

We have demonstrated that the G37V change is responsible for at least a 100-fold decrease in enzyme sensitivity to fenpicoxamid in single mutants, and a 20-fold decrease in double mutants. According to docking calculations, the molecular distance in the resistant cytochrome *b* between valine 37 and fenpicoxamid is less than 2 Å. This distance is barely longer than a C-C covalent bond (1.54 Å), suggesting that steric hindrance is likely. The mutation was also associated with a decrease in complex III activity, likely due to ubiquinone binding impairment, or to a negative impact of the mutation on electron transfer. Indeed, some Q_i_ site changes have been shown to impact negatively the electron transfer between the Q_o_ and Q_i_ sites and/or between the Q_o_ site and the Rieske iron-sulphur protein, hence reducing complex III activity (Cooley *et al*., 2005). A similar penalty has been described for the S34L substitution in *S. cerevisiae*, a mutation that triggers ametoctradin resistance in *P. viticola* (Mounkoro *et al*., 2019). The S34L change confers high resistance levels, but also a 2-fold decrease in enzyme activity in yeast. In *Z. tritici*, the 2-fold decrease in enzyme activity triggered by the G37V change had no detectable impact on *in vitro* growth, suggesting that complex III activity is not limiting in these conditions. A similar observation was made for SDH laboratory mutants which displayed decreased enzyme activity of up to 90% yet showed no visible growth defect *in vitro* (Scalliet *et al*., 2012). Some of the same mutations in these laboratory mutants were also identified in field isolates (Rehfus *et al*., 2018).

RFs for UK-2A against G37V strains were somewhat variable. Such variations may be explained by the presence of additional mutations with minor effects, which were likely acquired during experimental evolution. Differences in resistance levels to QiIs may also be due to heteroplasmy. This phenomenon has been reported for QoI resistance in various species like *Botrytis cinerea* (Hashimoto *et al*., 2015), *Venturia inaequalis* (Villani and Cox, 2014), or *Podosphaera xanthii* (Vielba-Fernández *et al*., 2018). The existence of a stable heteroplasmic state in *Z. tritici* has never been described but cannot be excluded.

We have also observed an interaction between G37V and the G143A changes, in the Q_i_ and Q_o_ binding sites of cytochrome *b*, respectively. The selection of G37V in parental strains containing G143A occurred less frequently than in the WT IPO323 strain, suggesting that G143A could have reduced G37V selection to some extent. Moreover, we highlighted the negative cross-resistance between QoIs and QiIs in single mutants resistant to one mode of action. An increased sensitivity to UK-2A was also apparent at the enzyme level in strains containing the G143A change (Table 3). The increased sensitivity may be a consequence of reduced enzyme efficiency associated with the G37V and G143A changes. Alternatively, a mutation at one site may increase inhibitor binding at the opposite site. The slightly lower RFs to both QoIs and QiIs in G37V+G143A double mutants also point to some form of interaction between the Q_o_ and Q_i_ sites.

### Consequences for fenpicoxamid use in the field

To our knowledge, this is the first study using directed evolution to predict resistance to QiIs. This confirms the interest of this approach to assess adaptation. Although laboratory mutant isolation studies cannot predict with certainty what will happen in the field, our results highlight the G37V change as a quite likely resistance mechanism since it was the only change detected and is a common locus for QiI resistance mutations in other organisms. The thorough characterization of G37V mutants showed that complex III activity was impaired in these isolates. Although their *in vitro* growth rate was similar to that of their ancestral strains, pathogenicity or fitness in the field could be affected, especially since some stages of the infection cycle require high energy production (Wood and Hollomon, 2003). In the case of QiI fungicides used to control oomycete diseases, the S34L change in the Q_i_ site of *P. viticola* causes resistance to ametoctradin. In a *S. cerevisiae* strain transformed with the *P. viticola* cytochrome *b* allele, this change produced a similar decrease (2-fold) in enzyme activity (Mounkoro *et al*., 2019) to that observed for G37V in *Z. tritici*. The S34L change remains at low frequency in French *P. viticola* populations, suggesting a fitness cost under field conditions. In contrast, the E203-VE-V204 insertion in the Q_i_ site of the same oomycete, associated with resistance to cyazofamid and amisulbrom, are thriving in French vineyards, suggesting a lower, or no fitness cost (Fontaine *et al*., 2019). Therefore, further risk assessment of the G37V mutants requires *in planta* studies.

The prediction of resistance in advance of its emergence in the field can enable sound and sustainable management of a new mode of action. Indeed, early prediction and detection of resistance increases the loss by chance of resistance alleles as they are maintained at low frequency in populations through an optimized management strategy. Molecular tools can be developed to allow early detection of anticipated mutants in real time (R4P-Network, 2016). In particular, molecular detection tools could allow to survey adjustments in usage of fenpicoxamid within a fine spatio-temporal scale. Detailed knowledge of the risk of resistance may also allow the smart and informed deployment of anti-resistance strategies, including mixtures, alternation or mosaic.

## EXPERIMENTAL PROCEDURES

### Isolates and media

Isolates used in this study were either collected in the field, or generated in the laboratory by experimental evolution or sexual reproduction between field isolates. Their genotypic variations in *MFS1* and cytochrome *b* are listed in Table 4.

**Table 4.** Origins and phenotypes/genotypes of *Z. tritici* isolates used in this study.

| Isolate | Cytochrome *b* | *MFS1* | Origin | Reference |
| --- | --- | --- | --- | --- |
| IPO323 | WT | WT | Reference isolate | Goodwin *et al.*, 2011 |
| 14-H-K3 | G143A | WT | Field isolate | Lab collection |
| 09-ASA-3a-PZ | G143A | MDR<sup>Type 1</sup> | Field isolate | Leroux and Walker, 2011 |
| 09-CB1 | G143A | MDR<sup>Type 1</sup> | Field isolate | Leroux and Walker, 2011 |
| 11183 | WT | MDR<sup>Type 1</sup> | Progeny isolate, lab cross | Lab collection |
| 35C1 | G143A | WT | Progeny isolate, lab cross | Lab collection |
| 35C3 | G143A | MDR<sup>Type 1</sup> | Progeny isolate, lab cross | Lab collection |
| 35C4 | G143A | MDR<sup>Type 1</sup> | Progeny isolate, lab cross | Lab collection |
| 35D16bis | G143A | WT | Progeny isolate, lab cross | Lab collection |
| 35F6 | G143A | WT | Progeny isolate, lab cross | Lab collection |
| 35F22 | WT | WT | Progeny isolate, lab cross | Lab collection |
| 35G5 | G143A | WT | Progeny isolate, lab cross | Lab collection |
| 37-15 | G143A | WT | Progeny isolate, lab cross | Lab collection |
| 37-16 | G143A | WT | Progeny isolate, lab cross | Lab collection |
| 37-19 | G143A | WT | Progeny isolate, lab cross | Lab collection |
| 37-22 | G143A | WT | Progeny isolate, lab cross | Lab collection |
| 37-41 | WT | MDR<sup>Type 1</sup> | Progeny isolate, lab cross | Lab collection |
| 37-59 | G143A | MDR<sup>Type 1</sup> | Progeny isolate, lab cross | Lab collection |
| W.F.2.44 | G37V | WT | Experimental evolution | This study |
| W.F.4.14 | G37V | WT | Experimental evolution | This study |
| W.F.5.10 | G37V | WT | Experimental evolution | This study |
| 716.3F.2.2 | G37V+G143A | WT | Experimental evolution | This study |
| 716.3F.2.3 | G37V+G143A | WT | Experimental evolution | This study |
| 716.3F.2.4 | G37V+G143A | WT | Experimental evolution | This study |
| 716.3F.3.1 | G37V+G143A | WT | Experimental evolution | This study |
| 716.3F.3.2 | G37V+G143A | WT | Experimental evolution | This study |
| 716.3F.4.1 | G37V+G143A | WT | Experimental evolution | This study |
| 716.3F.4.2 | G37V+G143A | WT | Experimental evolution | This study |
| 716.3F.4.3 | G37V+G143A | WT | Experimental evolution | This study |
| 716.3F.4.4 | G37V+G143A | WT | Experimental evolution | This study |
| 716.3F.4.5 | G37V+G143A | WT | Experimental evolution | This study |

**Table 5.** Maximum growth rates and statistical groups.

| Strain | Scal | SD | Group |
| --- | --- | --- | --- |
| IPO323 | 12,3255 | 0,0988 | a |
| W.F.5.10 | 12,3451 | 0,0979 | a |
| W.F.2.44 | 12,5582 | 0,0951 | a |
| W.F.4.14 | 12,3048 | 0,0974 | a |
| 37-16 | 15,214 | 0,147 | a |
| 716.3F.2.2 | 16,122 | 0,152 | b |
| 716.3F.3.2 | 15,616 | 0,141 | ab |
| 716.3F.4.4 | 15,776 | 0,143 | b |

**Table 6.** Strains’ maximum growth rate comparisons.

| Contrast | Estimate | SD | t.ratio | p-value |
| --- | --- | --- | --- | --- |
| IPO323 - W.F.5.10 | -0.0196 | 0.139 | -0.141 | 0.9990 |
| IPO323 - W.F.2.44 | -0.2327 | 0.137 | -1.697 | 0.3255 |
| IPO323 - W.F.4.14 | 0.0207 | 0.139 | 0.149 | 0.9988 |
| W.F.5.10 - W.F.2.44 | -0.2131 | 0.137 | -1.561 | 0.4013 |
| W.F.5.10 - W.F.4.14 | 0.0402 | 0.138 | 0.291 | 0.9914 |
| W.F.2.44 - W.F.4.14 | 0.2533 | 0.136 | 1.861 | 0.2455 |
| 37-16 - 716.3F.2.2 | -0.909 | 0.211 | -4.299 | 0.0001 |
| 37-16 - 716.3F.3.2 | -0.402 | 0.203 | -1.977 | 0.1973 |
| 37-16 - 716.3F.4.4 | -0.562 | 0.204 | -2.747 | 0.0309 |
| 716.3F.2.2 - 716.3F.3.2 | 0.507 | 0.208 | 2.440 | 0.0702 |
| 716.3F.2.2 - 716.3F.4.4 | 0.347 | 0.209 | 1.663 | 0.3440 |
| 716.3F.3.2 - 716.3F.4.4 | -0.160 | 0.201 | -0.796 | 0.8563 |

All isolates were stored in 25% glycerol at -80°C, and grown in liquid or on solid YPD (10 g/L yeast extract, 20 g/L peptone, 20 g/L glucose and 20 g/L agar if needed). Apical germ-tube length measurement establishing dose-response curves for tolnaftate sensitivity assays were performed on PG solid medium (10 g/L glucose, 2 g/L K_2_HPO_4_, 2 g/L KH_2_PO_4_ and 12.5 g/L agar).

### Fungicide sensitivity assays

#### Germ-tube elongation assay

*Z. tritici* isolates were grown at 18°C on YPD agar plates for 72h under light. Spore suspensions in sterile water were prepared from 3-day old cultures, and adjusted to 2.10^5^ spores/mL. 300 µL of spore suspensions were plated on PG medium containing increasing tolnaftate concentrations in ethanol (0.5 µg/mL final concentration), or control plates without fungicide containing 0.5 µg/mL final ethanol concentration. Plates were incubated for 48h in the dark at 17°C. After incubation, the length of apical germ-tubes from 10 spores was measured under the microscope using a micrometer (magnification x200). Tests were replicated independently three times.

#### Fungitoxicity assay in microtiter plates

*Z. tritici* isolates were grown at 18°C on YPD agar plates for 72h under light. Cells were then suspended in liquid YPD at 10^5^ spores/mL. Wells were filled with 1 µL of a concentrated fungicide solution in DMSO at 200 times the desired final concentration and 100 µL of YPD, then inoculated with 100 µL of the spore suspension. Final fungicide concentrations ranged between 0.0001 and 10 µg/mL for UK-2A and azoxystrobin, using a three-fold dilution series. Plates were incubated at 18°C for 72h. Growth was assessed in a NepheloStar plate reader (BMG Labtech, Cary, NC, USA), and EC_50_ values were determined by non-linear regression (curve-fit) using GraphPad Prism (GraphPad Software, La Jolla, CA, United States). EC_50_ values were the average of three technical replicates per assay and each assay was repeated at least twice.

### Selection and characterization of fenpicoxamid-resistant *Z. tritici* isolates

#### Experimental evolution

Ancestral isolates were either susceptible to QoIs and QiIs (IPO323), resistant to QoIs and susceptible to QiIs (35C1, 35D16bis, 35G5, 37-16, 37-19 and 37-22), MDR isolates susceptible to QoIs and QiIs (11183 and 37-41) or MDR isolates resistant to QoIs and susceptible to QiIs (35C4 and 37-59). Each experiment encompassed 3 or 4 independent lines for each ancestral isolate. 10^7^ spores were incubated at 17°C in 25 mL of liquid YPD containing 100 µg/mL propyl gallate and fenpicoxamid with shaking at 150 rpm in the dark for 7 days. Fenpicoxamid concentrations used were the MIC or 25MIC for each isolate, as established in preliminary experiments using the same experimental conditions. New selection cycles were initiated by transferring 0.5% of cells from the previous cycle to fresh medium. A concentration of 10^7^ cells at the start of each cycle was achieved by adding cells from an untreated control cell line. At the end of each cycle, 150 µL of each population were plated on fenpicoxamid-amended YPD at the selecting concentration, to detect resistant isolates. Experiments were stopped after eight selection cycles. Isolates were subcultured twice on fenpicoxamid-amended YPD before storage.

#### Characterization of fenpicoxamid-resistant isolates

Resistant isolates were grown in 25 mL of liquid YPD at 17°C with shaking at 150 rpm, in the dark for 3 days. OD at 560 nm was measured for each culture to standardize spore concentration. Spore suspensions were plated using three 10-fold serial dilutions on YPD agar with selecting compounds: 2 µg/mL fenpicoxamid, 0.5 µg/mL UK-2A, 2 µg/mL tolnaftate, 2.5 µg/mL azoxystrobin, 100 µg/mL propyl gallate, 100 µg/mL SHAM, and mixtures of fenpicoxamid and propyl gallate, or fenpicoxamid and SHAM, at the same concentrations. Plates were incubated at 17°C in the dark for 7 days before scoring. Scores ranged from 0 to 5 depending on growth intensity at each concentration.

#### Molecular characterization of resistant isolates

After total DNA extraction, *CYTb* PCR was performed using the primers Zt_Cytb_2_F (5’CCTGACTGGTATCATATTGTGT3’) and Zt_Cytb_1_R (5’TATATTACTAGGTTATTTTTCGTG3’). PCR conditions were an initial denaturation at 95°C for 3 min, followed by 35 cycles of 95°C for 20 s, 56°C for 20 s, 72°C for 2 min, and a final extension of 72°C for 5 min, giving a 1572 kb amplicon. Sequencing reactions with the primers Zt_Cytb_2_F and ZtCyto6 (5’ TAGGTTATTTTTCGTGTATAAAC 3’) were performed by Eurofins genomics (Ebersberg, Germany).

### Impact of target site mutations on complex III

#### Mitochondria extraction

*Z. tritici* mitochondria were prepared according to a protocol adapted from Lemaire and Dujardin, 2008 and Scalliet *et al*., 2012. Briefly, 10^7^ spores were incubated in 1 L of liquid YPD, at 17°C, in the dark with shaking at 150 rpm for 3 days. After centrifugation, the pellet was rinsed with sterile water, and immersed in liquid nitrogen before grinding. The resulting powder was resuspended in buffer A (0.7 M sorbitol, 50 mM Tris HCl pH 7.5, 0.2 mM EDTA pH 7.5). Mitochondria were prepared by differential centrifugations at 4°C: cell debris were eliminated at 3500 g for20 min then the supernatant was centrifuged at 13.000 g for 90 min. The pellet was resuspended in buffer B (buffer A with 0.5% BSA). Finally, cell debris were eliminated at 800 g for 5 min and mitochondria were pelleted at 15.000 g for 15 min and stored in buffer B (buffer A with 0.5% BSA) at –80°C.

#### Complex III activity

Decylubiquinol-cytochrome *c* reductase activities were determined at room temperature by measuring the reduction of cytochrome *c* (final concentration of 20 µM) at 550 nm *versus* 540 nm over a 2-min time-course. Mitochondria (final concentration to have a constant initial rate between 0.4 and 0.5) were added in 1 mL of 10 mM potassium phosphate pH 7, 0.01% (w/v) lauryl-maltoside and 1 mM KCN to inhibit complex IV activity. The reaction was initiated by adding decylubiquinol (final concentration of 40 µM) and the initial cytochrome *c* reduction rates were recorded. The measurements were performed in the presence of increasing concentration of inhibitors. The mid-point inhibition concentrations (I_50_) were estimated from the inhibitor titration plots. Each measurement was repeated at least twice and independent values were averaged.

#### Complex IV activity

The activity of complex IV was determined at 25°C by measuring oxygen consumption with an oxygen electrode. Complex IV activity was used to normalize complex III activity in the absence of a reliable method for quantifying cytochrome *bc*_1_ complex in *Z. tritici* mitochondrial extracts. Mitochondria were added to 1 mL of 50 mM potassium phosphate pH 7, 10 mM ascorbate and 50 μM N,N,N’,N’-tetramethyl-p-phenylenediamine at pH 7. The reaction was initiated by the addition of 50 μM cytochrome *c*. The initial oxygen consumption rates were measured at least 3 times and independent values were averaged.

#### In vitro kinetic growth assays

*Z. tritici* spores were suspended in liquid YPD at 10^5^ spores/mL. 200 µL of cell suspensions were placed in wells of 96-well microtiter plates and incubated at 18°C for 2h to allow spores to settle on the well surface. Plates were then incubated in a Cytation 3 instrument for 60h. Wells were imaged every 4 hours and confluence was calculated after Z-stacking (4 heights). Curves were modeled using the self-starting logistic model (sslogis) from the R software (The R Foundation for Statistical Computing), by the equation “Growth(Time) = Asym/(1 + exp((xmid – Time)/scal))”. “Asym” a numeric parameter representing the asymptote, “xmid” a numeric parameter representing the x value at the inflection point of the curve and “scal” is a numeric scale parameter on the input axis, here corresponding to the maximum growth rate. Maximum growth rates as Δconfluence/hour were determined. Statistical comparisons between strains were performed using the R package “emmeans”. Values of maximum growth rates and strain comparisons p-values are given in the following tables.

#### Molecular docking of UK-2A at the Z. tritici Q_i_ site

The *Z. tritici* homology model was built based on a template of the *S. cerevisiae* cytochrome *bc*_1_ complex (PDB: 1EZV) crystal structure (Hunte *et al*., 2000), which shares a sequence identity of 60% with the *Z. tritici* protein. The model was refined with the Amber14:EHT force field. UK-2A was then docked into the Q_i_ site of the model using induced fit docking. A total of 30 UK-2A binding poses were generated, from which the most likely binding pose was selected based on the docking energy score and knowledge of target site mutation effects in yeast (Young *et al*., 2018). All calculations were performed with the MOE software (Chemical Computing Group Inc., Montreal, QC, Canada).

## AKNOWLEDGEMENTS

Authors acknowledge the French National Association for Research and Technology (ANRT) for funding GF PhD through the CIFRE fellowship n°2017/1255. GF is also thankful to Agathe Ballu for her helpful participation in statistical analyses. All authors disclose they have no conflicts of interest relevant to this study.

